# A two-stage framework for neural processing of biological motion

**DOI:** 10.1101/2022.03.08.483404

**Authors:** João Valente Duarte, Rodolfo Abreu, Miguel Castelo-Branco

**Author notes:** **Corresponding author:** Miguel Castelo-Branco, Address: CIBIT – ICNAS, Instituto de Ciências Nucleares Aplicadas à Saúde, Pólo das Ciências da Saúde, Universidade de Coimbra, Azinhaga Santa Comba, Celas, 3000-548 | Coimbra, Portugal, |. Authors contributed equally to this work (João Valente Duarte, Rodolfo Abreu).

## Abstract

It remains to be understood how biological motion is hierarchically computed, from discrimination of local “life motion” animacy to global dynamic body perception. Here, we addressed this functional separation of the correlates of the perception of life motion, defined as characteristic for the local motion of parts of living beings, from perception of global motion of a body. We hypothesized that life motion processing can be isolated, by using a single dot motion perceptual decision paradigm featuring the biomechanical details of local realistic motion of a single joint. To ensure that we were indeed tackling processing of biological motion properties we used a discrimination instead of detection task. We discovered using representation similarity analysis that two key early dorsal and two ventral stream regions (visual motion selective hMT+ and V3A, extrastriate body area EBA and a region within fusiform gyrus FFG) showed robust and separable signals related to encoding of life motion and global motion. These signals reflected two independent processing stages, as revealed by representation dissimilarity analysis and deconvolution of fMRI responses to each motion pattern. This study showed that higher level pSTS encodes both classes of biological motion in a similar way, revealing a higher-level integrative stage, reflecting scale independent biological motion perception. Our results reveal a two-stage framework for neural computation of biological motion, with an independent contribution of dorsal and ventral regions for the initial stage.

## Introduction

Discrimination of biological motion, the movements of animate entities, is vital for survival. In humans, accurately perceiving dynamic bodily signals is crucial for nonverbal social cognition (Blakemore and Decety, 2001; Pavlova, 2012). Thus, the human visual system has evolved into an extremely sensitive mechanism to perceive biological motion, being capable of depicting human activity only via the motion of point lights at the main joints of a person, which are known as point-light displays – PLDs (Blake and Shiffrar, 2007; Johansson, 1973; Troje, 2013). Despite their apparent simplicity, human PLDs are characterized by a substantially higher degree of complexity than that of simple motion patterns typically used to study the visual system. With biological motion patterns, the visual system needs to integrate the individual dots, each with a different motion pattern - local motion, into the percept of a coherent body bearing multiple articulated joints - global motion. These two sources of information, “local animacy” (kinematics of individual dots) and “global” (structure from motion) may represent two processing stages.

Neuroimaging and neurophysiology studies have identified a network of areas that process biological motion. Relevant areas include the posterior superior temporal sulcus (pSTS) (Allison et al., 2000; Beauchamp et al., 2002; Bonda et al., 1996; Grossman et al., 2004, 2000; Grossman and Blake, 2002; Howard et al., 1996; Krakowski et al., 2011; Kriegstein and Giraud, 2004; Pavlova et al., 2004, 2007; Pelphrey et al., 2003; Puce and Perrett, 2003; Saygin et al., 2004), fusiform gyrus (FFG) (Grossman et al., 2004, 2000; Peelen et al., 2006; Vaina et al., 2001), inferior frontal gyrus (IFG) (Saygin, 2007; Saygin et al., 2004), extrastriate and fusiform body areas (EBA, FBA), (Bonda et al., 1996; Chang et al., 2018; Downing et al., 2001; Grosbras et al., 2012; Grossman and Blake, 2002; Jastorff and Orban, 2009; Peelen et al., 2006; Peuskens et al., 2005; Saygin, 2007; Spiridon et al., 2006; Thompson and Baccus, 2012; van Kemenade et al., 2012), insula (Freitag et al., 2008; Saygin et al., 2004; Sokolov et al., 2018), cerebellum (Sokolov et al., 2018, 2012), and low level visual areas, such as the human middle temporal complex (hMT+) and V3A (Chang et al., 2018; Grossman and Blake, 2002; Peelen et al., 2006). However, most experimental studies, including recent ultra-high-field fMRI (Pavlova et al., 2017) and connectivity studies (Dasgupta et al., 2017; Sokolov et al., 2018), do focus on global percepts of biological motion, in a way that only regions that respond to the structure-from-motion global aspects of biological motion are found. Therefore, the first stage’s neural correlates and mechanisms that allow an observer to interpret biological motion from the local motion of individual dots (Mather et al., 1992), or from point light stimuli deprived of any structural information (Troje and Westhoff, 2006), remain little understood.

Extensive behavioral data suggest that the visual system is remarkably sensitive to local kinematic information, and particularly responds to the gravitational acceleration of the feet (Chang and Troje, 2009a; Troje and Westhoff, 2006). This may be enabled by using strictly defined operational definitions, such as “life motion”, which reflects aspects that are exclusive of the motion of living beings (Troje, 2013). The notion of a “life detector” was suggested in an experiment where inverting the trajectory of feet motion entirely disrupted the ability to determine the facing direction of biological motion PLDs (Troje and Westhoff, 2006). This sensitivity to the invariants in foot motion and its independence of shapes (Chang and Troje, 2009a, 2009b; Hirai et al., 2011b; Jiang and He, 2007; Wang et al., 2010) might be present in human neonates (Bardi et al., 2011; Simion et al., 2008) and non-human species as well (Vallortigara and Regolin, 2006), which points to an evolutionarily old origin. This concept introduces an early (first stage) processing step in biological motion perception and raises the question of which are the neural correlates underlying life motion information processing prior to integration into a globally coherent articulated biological body moving structure. A recent fMRI study of the neural representations of articulated body shape and local kinematics in biological motion suggests the involvement of a broad set of cortical visual regions and the ventral lateral nucleus of the thalamus (VLN) (Chang et al., 2018). However, they focused on global and distributed multiple local cues, and the contribution of local life motion from an isolated source of information (e.g., one foot) and its role in a two-stage processing system remains to be discovered. Thus, local foot motion, which is a component of a living body motion, may represent an appropriate approach to isolate the first stage of biological motion processing.

We therefore developed an event-related fMRI experimental design to study the neural correlates of the two-stage system of biological motion processing. We used the concept of life motion - featuring a first processing stage - based on a single moving dot embedding the kinematic properties of local foot motion, as well as structural global motion perception - featuring both first and second processing stages - based on PLDs. Since no information about body structure or articulate motion was present during single foot motion, we could isolate more low-level (first stage) aspects of biological motion processing from aspects related to social perception of animated shapes (second stage). We then asked whether local biological (life motion) and global biological motion is processed in the same network or if the two processing stages rely on different areas. We did not intend to directly look at local vs. global contrasts, rather to understand the biological motion network processing in two different contexts (local and global) in discrimination tasks. Furthermore, since local and global motion displays differ greatly in low-level visual properties, we used visually-matched control conditions. We showed the same displays of dots (foot or body) with manipulated motion profiles, eliciting very similar perceptions of (local and global) biological motion but physically deviating from veridical biological motion. We used, for the first time, as to our knowledge, event-related deconvolution GLM analysis and representational similarity analysis (RSA) to investigate the two-stage framework for neural processing of biological motion. We hypothesize that the first stage is involved in local and global motion, while the second stage is involved mainly in global motion processing. We also manipulated the stimuli in terms of the constraints imposed by gravity and inertial forces of each dot, resulting in slightly smoothed acceleration/deceleration patterns of motion, to study the behavior of the two stages when there is degradation of veridical biological motion patterns.

In sum, in this fMRI study we aimed to address the neural basis the two stages of biological motion processing: life motion perception, using single dot stimuli that only contained local properties, and global biological motion perception. We found that the first stage relies on early visual areas hMT+ and V3A and higher-level areas FFG and EBA, which seem to discriminate local and global motion. The second stage seems to rely on pSTS, which is indifferent to the subtype of biological motion.

## Materials and Methods

### Participants

Seventeen healthy participants (mean age: 28 ± 6 years; 9 males) were recruited. All participants had normal or corrected-to-normal vision, no history of neurological or psychiatric disorders, and all were right-handed, as confirmed by a handedness questionnaire (Oldfield, 1971). The study was approved by the Ethics Commission of the Faculty of Medicine of the University of Coimbra and was conducted in accordance with the declaration of Helsinki. The study complied with the safety guidelines for magnetic resonance imaging (MRI) research in humans. All participants provided written informed consent to participate in the study.

### Experimental setup

MRI scanning was performed on a 3T Siemens Magnetom Prisma Fit scanner (Siemens, Erlangen, Germany) using a 64-channel RF receiver head coil, at the Institute of Nuclear Sciences Applied to Health (ICNAS), University of Coimbra, Portugal.

Visual stimulation was implemented using the Psychophysics Toolbox version 3 (Brainard, 1997; Pelli, 1997) and the BioMotion toolbox (van Boxtel and Lu, 2013) in MATLAB (The MathWorks, Inc., Natick, MA-USA). Stimuli were shown inside the MR scanner bore by means of an LCD screen (Avotec Real Eye Silent Vision 6011, Stuart, FL-USA) with resolution 1920 × 1080 and refresh rate 100 Hz, located at the rear of the magnet approximately ∼156 cm away from the participants’ eyes. Participants viewed the screen through a 45 deg tilted mirror mounted above their eyes. Behavioral responses were collected using a fiber-optical MRI-compatible response pad, Lumina LS-PAIR (Cedrus Lumina LP-400, LU400 PAIR, Cedrus Corporation, San Pedro, CA-USA).

### hMT+/V3A functional localizer stimulus

A functional localizer (single-run) was used to find the subject-specific location of hMT+ and V3A. For this purpose, visual stimulation alternated between moving and stationary dot patterns in the center of the screen with 600 high contrast white dots on a black background. Dots traveled toward and away from a central fixation cross at a constant speed (5 deg/sec) for 18 seconds, followed by a 10 second pattern of static dots. This cycle was repeated 12 times. Participants were instructed to watch passively but attentively. The hMT+/V3A were then identified as the voxels in anatomically valid locations that responded significantly higher to moving dots than to static dot fields (Huk et al., 2002).

### Biological motion stimuli

Biological motion stimuli were built based on human motion capture data collected at 60 Hz, comprising 12 point-lights placed at the main joints of a male walker facing rightwards. Stimulus duration is 0.75 seconds (45 samples), spanning one gait cycle. Overall translation was subtracted so that the point-light walker is walking on the spot. These data are part of the distribution of the BioMotion toolbox (van Boxtel and Lu, 2013). The point-lights were white dots displayed on a black background.

The walker in its original form contains both full structure from motion information (through the presence of the familiar body shape) and kinematic information (as carried by horizontal and vertical asymmetries, such as acceleration). To tease apart the contributions of these differing types of information, we presented the walker in its original form (GLOBAL motion) and only the right foot of the walker (LOCAL motion).

### Manipulation of biological motion patterns

We derived three variations of each stimulus type (GLOBAL and LOCAL), manipulating the kinematic information (temporal component) contained in the individual local trajectories in terms of the tangential local velocity (*ν*), that is, the velocity along the translational pattern, while keeping constant the average speed and length of the gait cycle. The velocity of the translation pattern was smoothed using a moving average (smoothed average: SM) with 9 (SM1), 23 (SM2) and 36 (SM3) samples; higher number of samples yields smoother translational patterns, and therefore, farthest from the true translational pattern (true biological motion, BM). For each smoothed velocity pattern, a new set of 3D coordinates (translation, elevation and sagittal) was computed. In this way, all four versions (BM, SM1, SM2 and SM3) of each stimulus type (GLOBAL and LOCAL) share the same spatial properties, but comprise different temporal characteristics, with the toe-off and heel-strike being the same for each step. A total of 8 different conditions were then defined, four for each stimulus type: GLOBAL-BM, GLOBAL-SM1, GLOBAL-SM2, GLOBAL-SM3, LOCAL-BM, LOCAL-SM1, LOCAL-SM2 and LOCAL-SM3. The right foot point-light was the same in GLOBAL and LOCAL corresponding stimuli.

### Biological motion discrimination task

The biological motion discrimination task was designed under a mixed block/event-related framework (Petersen and Dubis, 2012), coupled with a two-alternative forced choice decision (2AFC) task (“which of two consecutive stimuli is closest to true biological motion?”), which is depicted in Figure 1. Notably, the stimuli were reported by participants to be very similar within each motion type (local or global), thus keeping the discrimination task within a scenario of biological motion visual stimulation, i.e., all stimuli looked biological to a certain degree. This ensured that the task was always engaging for the participants and avoided habituation effects.

**Figure 1.**
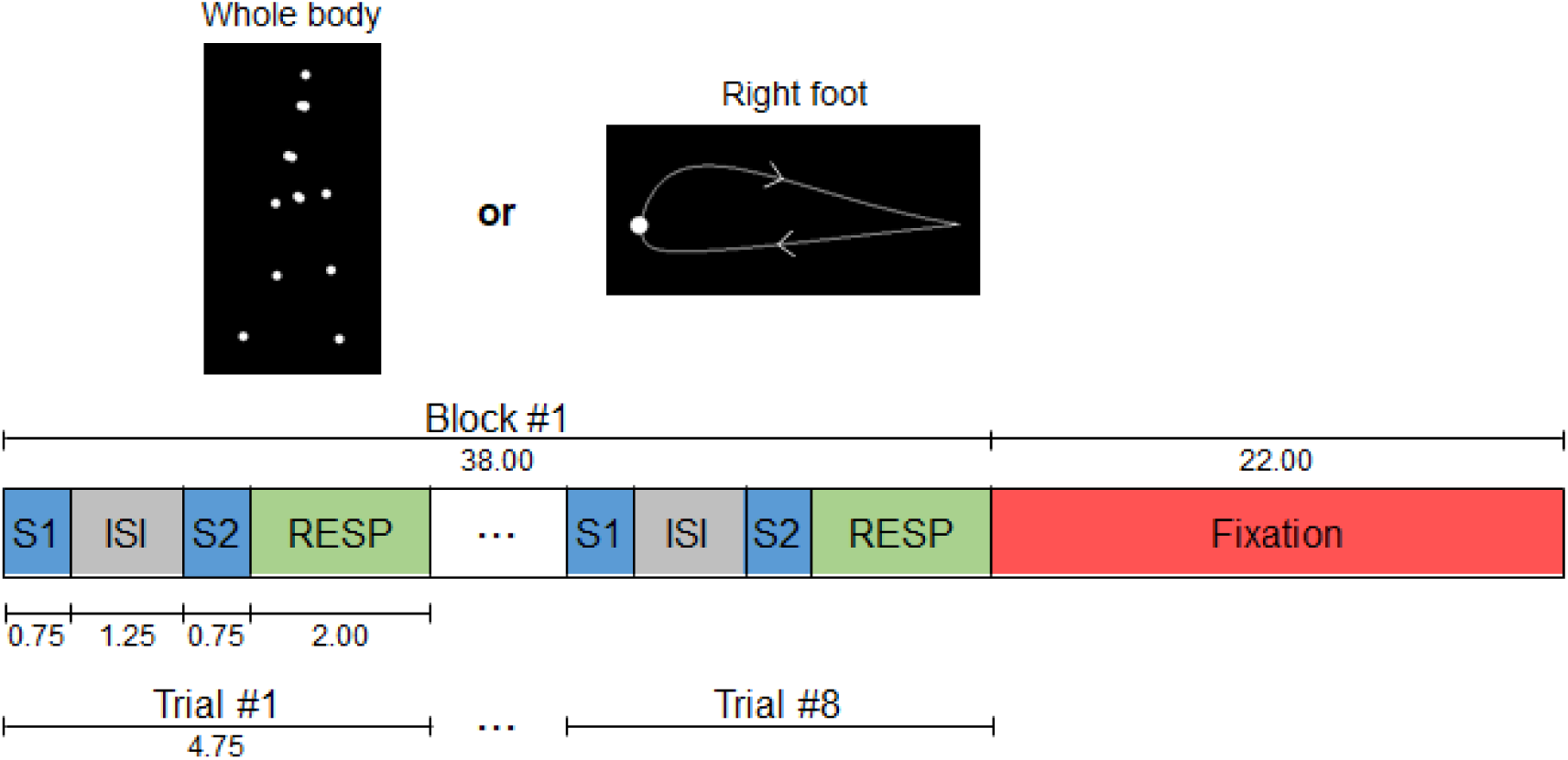
Schematic representation of the 2AFC biological motion discrimination task. The duration of each period is indicated in seconds. In each run, within each of four task blocks, there were eight trials of the discrimination task. S1: first stimulus; ISI: inter stimulus interval; S2: second stimulus; RESP: response period.

Participants were submitted to four consecutive runs, each consisting of 4 stimulation blocks of the conditions, 2 for each stimuli type. The blocks of global and local were interleaved and the starting block type was alternated across runs and participants. Each stimulation block lasted 38 seconds and comprised eight trials of 4.75 seconds; in each trial, two motion patterns of 0.75 seconds were shown, separated by a 1.25 seconds’ black screen period. The participants were asked at each trial to discriminate, via button press, which of the two patterns was the closest to true biological motion (the veridical biological motion, if presented, or the less smoothed gait cycle), and give their answer during a 2 second response period marked by a white cross following the second motion stimulus; no feedback on the response correctness was provided. Each stimulus was pseudo-randomly presented 20 times in each run, 80 times in total across the four runs. Each block was followed by a 22 second baseline period, during which a red fixation cross was displayed at the center of the screen.

### fMRI data acquisition

For each participant, the scanning session consisted of i) one T_1_-weighted (T_1_w) 3D anatomical magnetization-prepared rapid gradient echo (MPRAGE) pulse sequence (acquisition time 6 min 03 sec; repetition time (TR) = 2530 msec; echo time (TE) = 3.5 msec; inversion time (TI) = 1100 msec; flip angle 7°; 192 single shot slices with voxel size 1 × 1 × 1 mm^3^; field of view (FOV) = 256 × 256 mm^2^); ii) one functional run for the localization of hMT+/V3A using a T2*-weighted gradient echo (GE) echoplanar imaging (EPI) sequence with measurement of blood-oxygen-level-dependent (BOLD) signal (time of acquisition = 6m22sec; TR 1000 msec; TE = 30.2 msec; flip angle 68°, 51 interleaved slices with voxel size 2.5 × 2.5 × 2.5 mm^3^; multi-band acceleration factor = 3; FOV = 195 × 195 mm^2^); and iii) four functional runs for the biological motion discrimination task using the same sequence parameters used in the localizer run, except with acquisition time = 10m36sec, and an event-related design stimulation paradigm to measure BOLD signal changes in response to GLOBAL and LOCAL stimuli. Prior to each functional run, a short (10 volumes) GE-EPI acquisition with reversed phase encoding direction was performed, for image distortion correction.

### Data preprocessing

#### fMRI data cleanup

The first 10 s of data were discarded to allow the fMRI signal to reach steady-state, and non-brain tissue was removed using FSL’s tool BET (Smith, 2002). Subsequently, slice timing and motion correction were performed using FSL’s tool MCFLIRT (Jenkinson et al., 2002), followed by a B_0_-unwarping step with FSL’s tool TOPUP (Andersson et al., 2003) using the reversed-phase encoding acquisition, to reduce EPI distortions. Then, nuisance fluctuations were removed and a high-pass temporal filtering with a cut-off period of 100 s was applied; no spatial smoothing was applied. The nuisance fluctuations (including physiological noise) were modeled by linear regression using the following regressors (Abreu et al., 2017): 1) quasi-periodic fluctuations related to cardiac and respiratory cycles were modeled by a fourth order Fourier series using RETROICOR (Glover et al., 2000); 2) aperiodic fluctuations associated with changes in the heart rate as well as in the depth and rate of respiration were modeled by convolution with the respective impulse response functions (Chang et al., 2009); 3) the average BOLD fluctuations in white matter (WM) and cerebrospinal fluid (CSF); 4) the six motion parameters estimated by MCFLIRT; and 5) scan nulling regressors (motion scrubbing) associated with volumes acquired during periods of large head motion – motion spikes (Soares et al., 2021); these were determined using the FSL’s utility *fsl_motion_outliers*, whereby the DVARS metric proposed in (Power et al., 2012) is first computed, and then thresholded at the 75th percentile plus 1.5 times the inter-quartile range. For each participant, the functional images were linearly co-registered with the respective T_1_-weighted structural images using FSL’s tool FLIRT, and subsequently with the Montreal Neurological Institute (MNI) (Collins et al., 1994) template, using non-linear transformations estimated with the FSL’s tool FNIRT FNIRT (Jenkinson et al., 2002; Jenkinson and Smith, 2001).

### Data analysis

Imaging data analysis was carried out using BrainVoyager 21.2 (Brain Innovation, Maastricht, The Netherlands), and custom scripts in MATLAB R2020a (The Mathworks, Inc., Natick, MA-USA).

#### Behavioral analysis

We calculated the percentage of correct discrimination between each condition of motion and all the other conditions of the same type of motion, global and local. We then tested, with a two-way ANOVA with repeated measures with TYPE of stimuli (global and local) and SMOOTH level (BM, SM1, SM2 and SM3) as factors, the interaction of both in mean correct responses the simple main effects of each factor. In case of significant simple main effect, we tested the pairwise differences with Bonferroni correction for multiple comparisons. Significant differences were considered at *p*-value < 0.05.

#### ROI definition

The hMT+ and V3A regions-of-interest (ROIs) were functionally localized in every participant with an individual general linear model (GLM) analysis of the localizer run by contrasting the periods of moving dots that traveled toward and away from fixation with the periods of static dot fields (Huk et al., 2002). The individual statistical maps were limited at threshold value of *p-value* = 0.05, with Bonferroni correction. The correspondence between activated clusters and motion-sensitive areas hMT+ and V3A was confirmed by comparing their location with previous reports of functional maps obtained with retinotopic and flow-field mapping experiments (Castelo-Branco et al., 2002; Duarte et al., 2017; Tootell et al., 1997), as well as studies of biological motion processing (Chang et al., 2018; Hillebrandt et al., 2014; Kim et al., 2011; Sokolov et al., 2018). For further analysis, the left and right hMT+ and left and right V3A ROIs were used combined as two single ROIs, which we designate by bilateral hMT+ and bilateral V3A, or just hMT+ and V3A for sake of simplicity.

We included four additional extrastriate regions that have been previously implicated for biological motion perception (Chang et al., 2018). Each was defined as a spherical ROI (5 mm radius) centered on MNI coordinates of: [±58, -46, 6] for the posterior superior temporal sulcus (pSTS), [±37, 10, 28] for the inferior frontal gyrus (IFG), [left - 46, -75, -4; right 47, -71, -4] for the extrastriate body area (EBA), and [left -38, -38, -27; right 43, -43, -28] for the fusiform body area (FBA). A subcortical ROI corresponding to the ventral lateral nucleus (VLN) of the thalamus was also defined at coordinates [left - 16, -9, 8; right 14, -9, 8]. Three additional ROIs recently found to be functionally synchronized during different stages of biological motion processing were considered (Pavlova et al., 2017; Sokolov et al., 2018); these were located at [±42, -56, -14] for the fusiform gyrus (FFG), [±36, 24, 2] for the anterior insula (aINS), and [±36, -54, -28] for the lateral cerebellar lobule Crus I (Crus).

#### Neural responses global and local biological motion

Statistical analysis was performed using random effects (RFX) in order to generalize findings to the population level (Penny et al., 2003), from the concatenation of the four runs of each participant. We built predictors of interest for each stimulation event, one for each of the 8 combinations of motion type (GLOBAL and LOCAL) and smoothness level.

### Standard ROI-GLM

We performed a standard ROI-GLM analysis, with convolution of each predictor with the canonical hemodynamic response function (HRF), within subject-specific hMT+ and V3A, as well as within the other ROIs defined from MNI coordinates. We extracted the beta values per condition, representing the percentage of BOLD signal change relative to baseline, for each subject. We then compared the amplitude of response for each condition. We performed a three-way ANOVA with repeated measures to test the interaction effect of the factors ROI, TYPE of stimuli (global and local) and SMOOTH level (BM, SM1, SM2 and SM3), with follow-up Bonferroni-corrected two-way or one-way ANOVAs and post-hoc pairwise comparisons of each factor’s effect in brain responses. Whenever the assumption of sphericity was not met, we used the Greenhouse-Geisser correction in statistical reporting. Significant differences were considered at *p*-value < 0.05.

### Event-related ROI-deconvolution GLM

An additional strategy, novel in fMRI studies of biological motion, was also used to deal with the overlap of responses to different conditions. We employed a GLM deconvolution analysis, relying on a finite impulse response (FIR) model, within each ROI to estimate the profile of BOLD responses for each condition. The FIR model makes no assumption about the shape of the hemodynamic response and is useful in cases where differences in time courses may exist across conditions. This allows a more flexible fitting of the model and allows to compare conditions on a single data point basis (see, e.g., (Glover, 1999) for a detailed explanation). We then contrasted with paired t-tests the peak response (between 3 to 6 seconds) to biological motion against the response to artificial patterns of motion, separately for each stimulus type (global or local).

### Representational Similarity Analysis (RSA)

Representational similarity analysis (RSA) was used to analyze the similarity between fMRI responses evoked by the stimulation conditions (Kriegeskorte et al., 2008). For each ROI a representational distance (or dissimilarity) matrix (RDM) was defined by first computing the spatial pairwise Pearson correlation coefficient *r* between the BOLD response across voxels within each ROI; *r* ranges from -1.0 (perfect anti-correlation) over 0.0 (no correlation) to +1.0 (perfect correlation). The distance measure *d* is then defined as: *d* = 1 - *r*, and thus ranges from 0.0 (minimum distance, *r* = 1) to 2.0 (maximum distance, *r* = -1), with 1.0 representing the no correlation case (*r* = 0).

Additionally, this RSA procedure was also applied to the event-related responses estimated with the deconvolution approach, at each ROI and for each condition, with the purpose of disentangling overlapping responses to different conditions.

Finally, a multi-dimensional scaling (MDS) of the responses within each ROI was performed, whereby the triangular part of each RDM (computed with both approaches) is projected onto 2-dimensional and 3-dimensional spaces that maximally satisfy the pairwise distance of each condition to all other conditions. The resulting MDS plots (one 2D and one 3D per RDM) better illustrate the similarity structure coded in the RDMs. Furthermore, we applied second-level RSA and MDS analyses, considering the similarity of the responses of all ROI pairs, independently of stimulus type. This approach allows to infer which ROIs have more similar patterns of responses.

## Results

### Behavioral analysis

The percentage of correct discrimination between different versions (smooth) of each stimulus’ type is presented in Figure 3, separated for the global and local types of stimuli. There were no outliers, as assessed by examination of studentized residuals for values greater than ±3 (standard deviations). The proportion of correct discriminations was normally distributed for each condition, as assessed by Shapiro-Wilk’s test of normality on the studentized residuals (*p* > 0.05). According to the Mauchly’s test, the assumption of sphericity was met for the two-way interaction of TYPE of stimuli (global or local) and SMOOTH(ness) level (BM, SM1, SM2, and SM3). Thus, using a two-way ANOVA with repeated measures we observed a statistically significant two-way interaction between the TYPE of stimuli and SMOOTH level on behavioral performance, *F*(3,45) = 3.225, *p* = 0.031, η^2^ = 0.177. Subsequently, the simple main effect of TYPE of stimuli showed that the discrimination performance in trials of global type was statistically significantly higher compared to that in trials of local type of stimuli for all SMOOTH levels (all *p* < 0.004), as expected from the nature of cues. In any case, task performance levels were in both conditions indicative of a high level of ambiguity (see Figure 2). The simple main effect of SMOOTH in each stimuli TYPE revealed that mean discrimination accuracy was statistically significantly different over different SMOOTH levels only when the stimuli to discriminate were of the local type, *F*(3,45) = 6.460, *p* = 0.001, η^2^ = 0.301. Post hoc pairwise comparisons with Bonferroni correction for multiple comparisons showed that the proportion of correct responses was lower for the BM stimuli when compared to any other SMOOTH level (*p*_SM1_ = 0.044, *p*_SM2_ = 0.024, *p*_SM3_ = 0.045), showing that the perceptual decision task was indeed challenging.

**Table 1:**
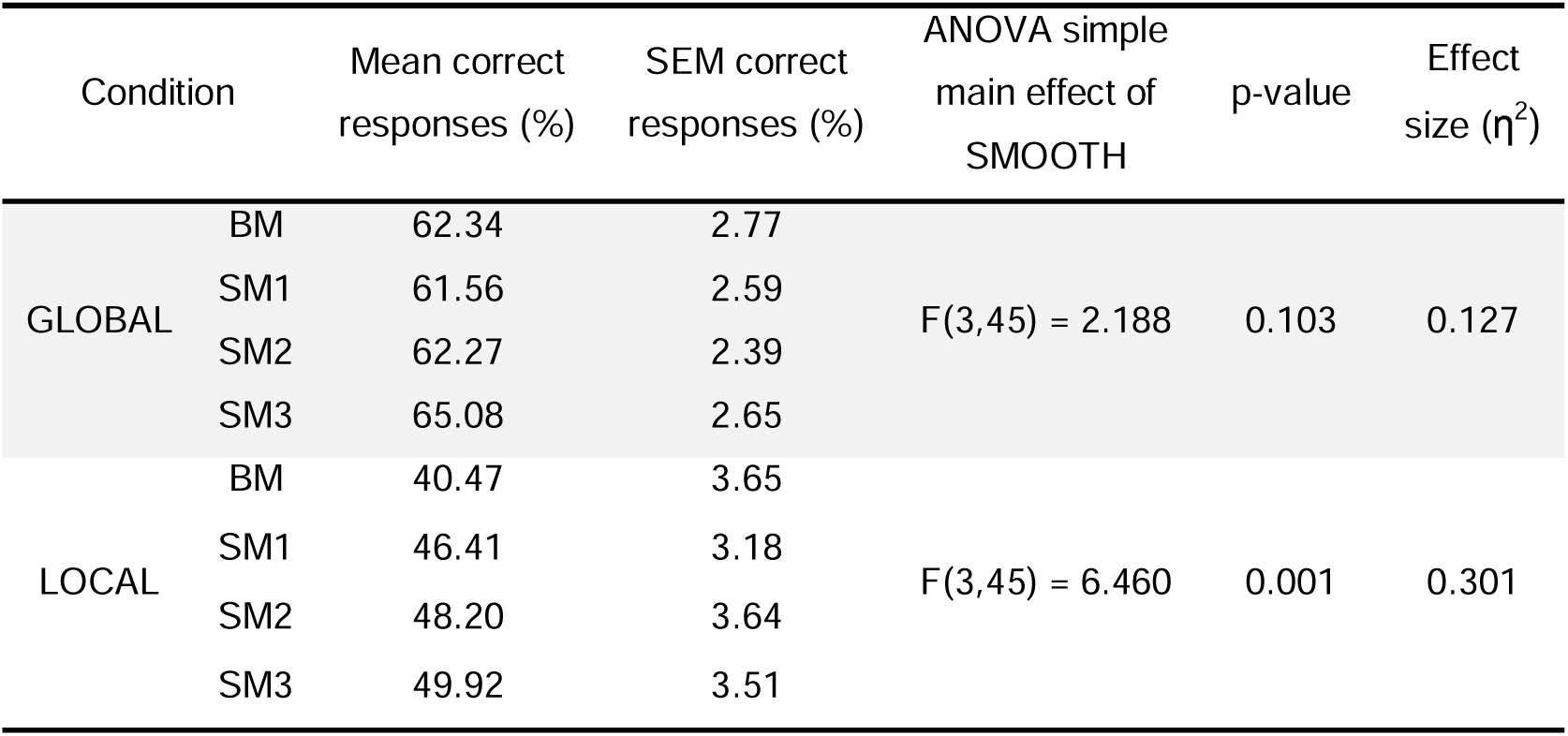
Proportion of correct responses to the discrimination between each stimulus *and all the other stimuli of the same type (global or local), performance being expectedly better for the largest stimulus differences*. There was a statistically significant two-way interaction between the TYPE of stimuli (global or local) and SMOOTH level (BM, SM1, SM2 and SM3) on behavioral performance, *F*(3,45) = 3.225, *p* = 0.031, η^2^ = 0.177.

| Condition | | Mean correct responses (%) | SEM correct responses (%) | ANOVA simple main effect of SMOOTH | p-value | Effect size ($\eta^2$) |
| --- | --- | --- | --- | --- | --- | --- |
| GLOBAL | BM | 62.34 | 2.77 | $F(3,45) = 2.188$ | 0.103 | 0.127 |
|  | SM1 | 61.56 | 2.59 |  |  |  |
|  | SM2 | 62.27 | 2.39 |  |  |  |
|  | SM3 | 65.08 | 2.65 |  |  |  |
| LOCAL | BM | 40.47 | 3.65 | $F(3,45) = 6.460$ | 0.001 | 0.301 |
|  | SM1 | 46.41 | 3.18 |  |  |  |
|  | SM2 | 48.20 | 3.64 |  |  |  |
|  | SM3 | 49.92 | 3.51 |  |  |  |

**Figure 2.**
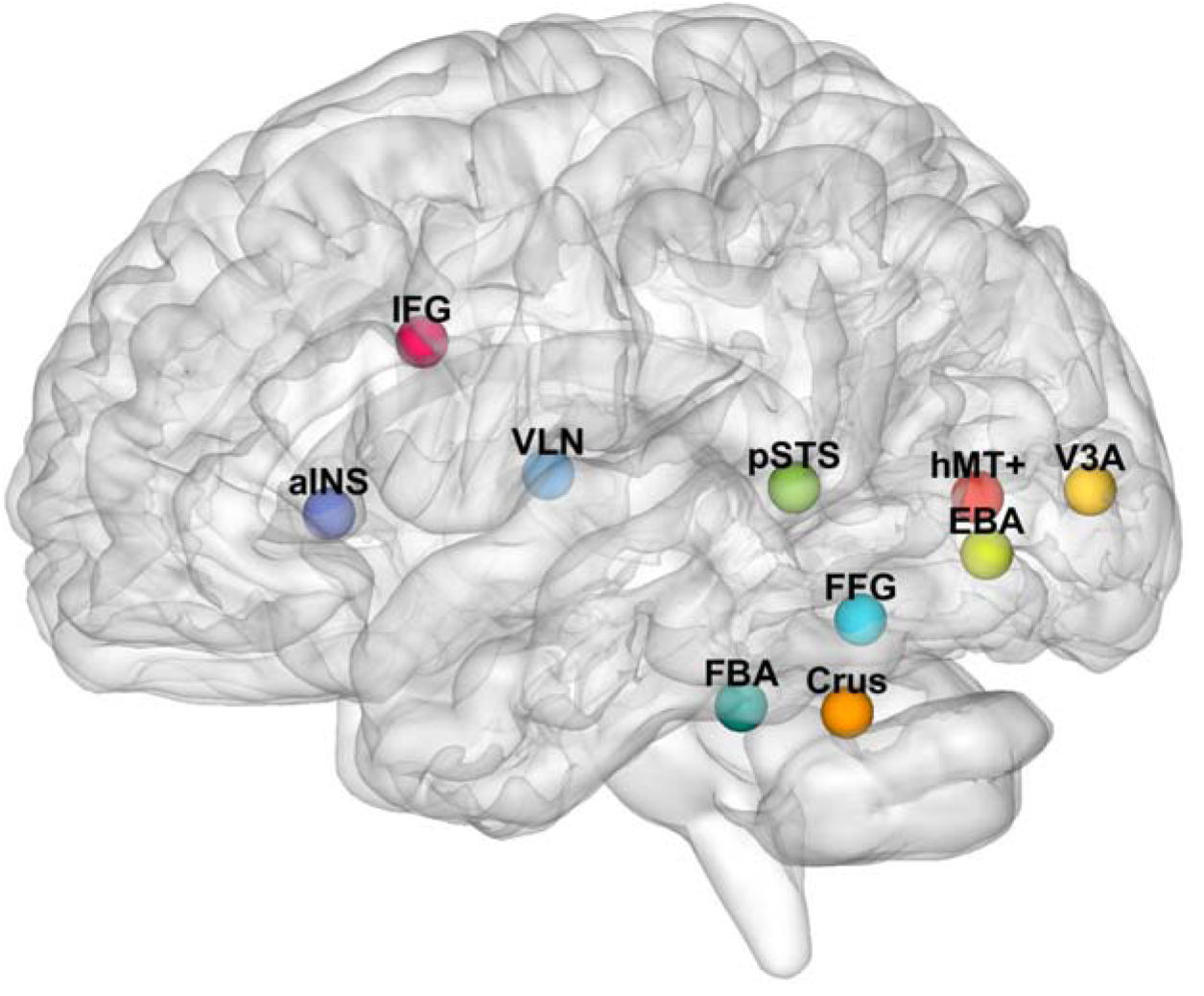
Location of the ROIs used in imaging analysis are overlaid on a 3D brain template. aINS: anterior insula; IFG: inferior frontal gyrus; VLN: ventral lateral nucleus of the thalamus; pSTS: posterior part of superior temporal sulcus; FBA: fusiform body area; FFG: fusiform gyrus; Crus: lateral cerebellar lobule Crus I; EBA: extrastriate body area; hMT+: human middle temporal complex.

**Figure 3.**
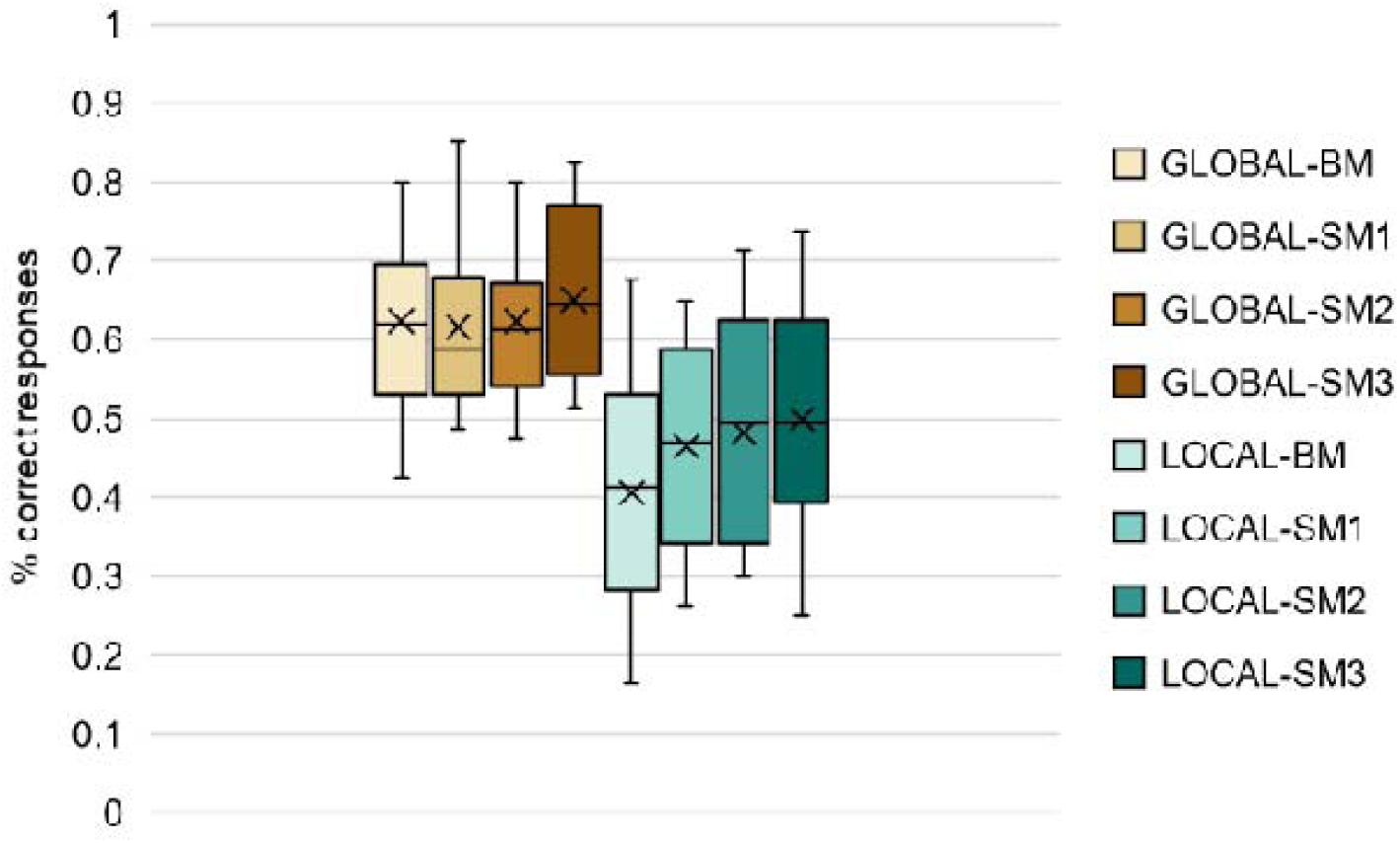
Behavioral (N=17) discrimination accuracies for trials presenting each type of stimulus (*against any other stimulus of the same type, global or local*). Boxes represent the interquartile range (IQR = Q3 – Q1), the horizontal line represents the median, and the crosses represent the average values. Whiskers (vertical lines) extend from the ends of each box to the minimum value and maximum value. BM = biological motion; SM = smoothed motion (the larger the number, the larger the violation of biological motion by smoothing of acceleration patterns; SM1 is the stimulus most similar to biological motion – see Methods). Asterisks indicate pairwise significant differences, revealed with Bonferroni-corrected post hoc tests.

### Imaging results

#### Standard ROI-GLM

The percentage of BOLD signal change in each ROI in response to each stimulus condition, compared to baseline, is presented in Figure 4, separated for the stimuli presenting global and local motion patterns. There were no outliers in the data, as assessed by inspection of boxplots. Beta values (% BOLD signal change) were normally distributed, as assessed by Shapiro-Wilk’s test of normality (*p* > 0.05). We did not observe a statistically significant three-way interaction between the ROI, the TYPE of stimuli (global or local), and the level of SMOOTH on brain responses, *F*(8.656,138.497) = 0.229, *p* = 0.077. The two-way ANOVA indicated a significant interaction between ROI and TYPE, *F*(4.219, 67.497) = 33.422, *p* = 9.473 × 10^−16^, η^2^ = 0.676, indicating that the differences in brain responses to different types of motion are dependent on the ROI considered.

**Figure 4:**
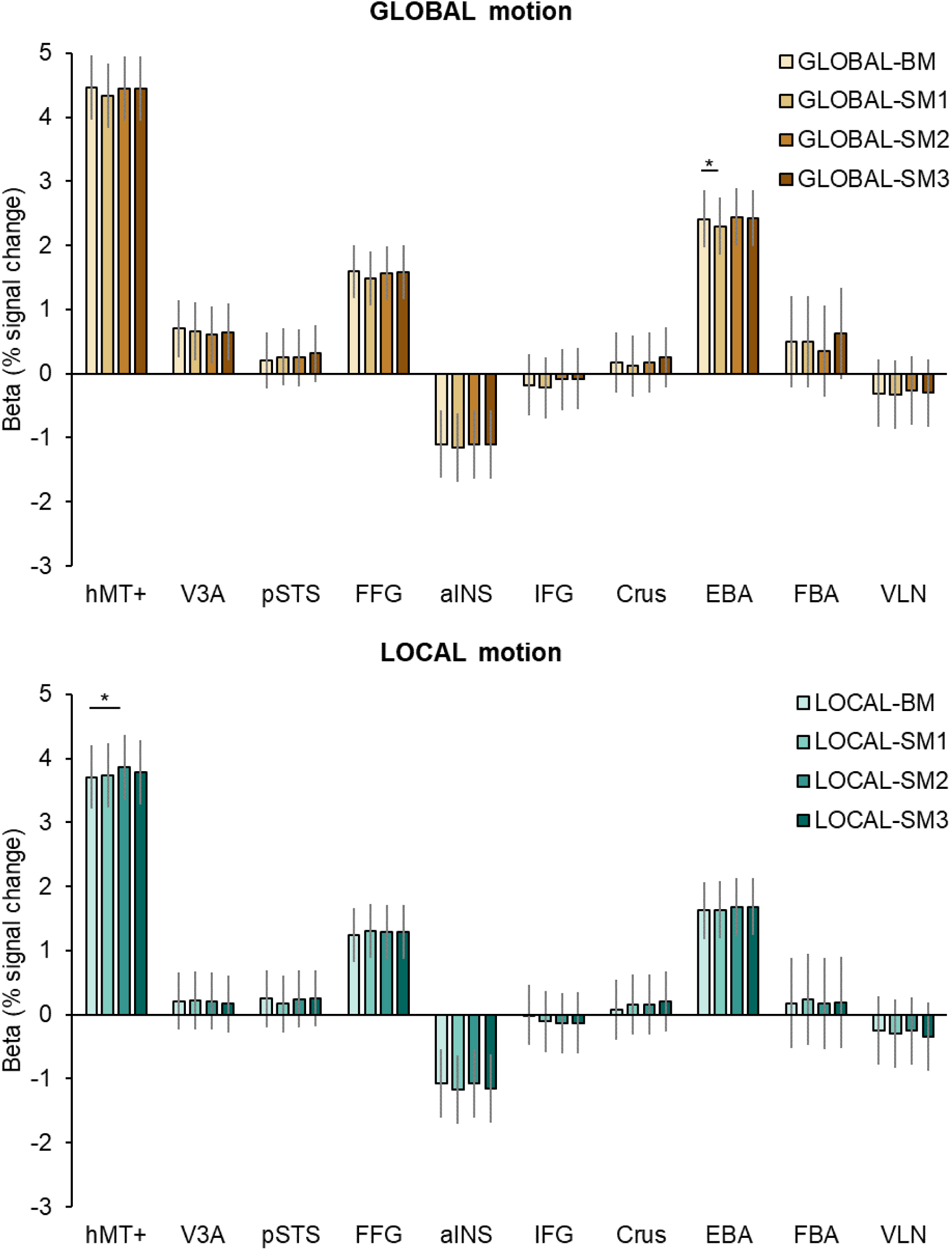
The percentage of BOLD signal change computed with standard GLM analysis of BOLD signal in each ROI, in response to each stimulus condition compared to baseline, for both global and local motion types.

A significant main effect of ROI on percent BOLD signal change was found, F(3.639,58.222) = 36.199, *p* = 1.460 × 10^−14^, η^2^ = 0.693, and follow-up comparisons revealed that the percentage of BOLD signal change was overall higher in hMT+ than in all other ROIs (all comparisons involving hMT+, *p* < 0.002). The signal in pSTS, commonly associated with biological context processing, was significantly lower than in FFG (*p* = 4.06 × 10^−4^) and EBA (*p* = 1.3 × 10^−5^). Additionally, BOLD signal change in EBA was lower than in hMT+ but significantly higher than in all other ROIs (all *p* < 0.001) except FFG and V3A. Similarly, FFG showed significantly higher responses than aINS (*p* = 1 × 10^−6^), IFG (*p* = 3.071 × 10^−3^), Crus (*p* = 0.025), FBA (*p* = 6.279 × 10^−3^) and VLN (*p* = 1.3 × 10^−5^). Notably, BOLD signal change in V3A was not significantly lower than in any other ROI. Regarding the stimuli TYPE, beta values were overall higher for the stimuli presenting global motion, which was confirmed by the significant main effect of type, F(1,16) = 101.469, *p* = 2.482 × 10^−8^, η^2^ = 0.864.

We also observed a trend for a two-way interaction effect of ROI x SMOOTH, *F*(8.775,140.395) = 1.764, *p* = 0.082], suggesting that the response to different levels of smoothing of the motion pattern is also dependent of the ROI in question. Thus, we followed these results with separate two-way ANOVAs in each ROI and follow-up one-way ANOVAs and post-hoc pairwise comparisons of the effect of each SMOOTH level in case of significant interaction between TYPE and SMOOTH. The analyses indicated that the pattern of responses across ROIs was slightly different for global motion stimuli compared to local motion stimuli. Signals in hMT+ were higher for global motion, revealed by the main effect of motion TYPE, *F*(1,16) = 66.216, *p* = 4.443 × 10^−7^, η^2^ = 0.805. The main effect of motion TYPE was also significant in V3A [*F*(1,16) = 46.600, *p* = 4 × 10^−6^, η^2^ = 0.744], in FFG [*F*(1,16) = 29.883, *p* = 5.2 × 10^−5^, η^2^ = 0.651], in EBA [*F*(1,16) = 58.455, *p* = 9.933 × 10^−7^, η^2^ = 0.785], and in FBA [*F*(1,16) = 34.706, *p* = 2.3 × 10^−5^, η^2^ = 0.684]. However, the main effect of SMOOTH was only significant in hMT+ [*F*(3,48) = 3.459, *p* = 0.023, η^2^ = 0.178] and the cerebellar lobule Crus I [*F*(3,48) = 0.2979, *p* = 0.041, η^2^ = 0.157].

#### Event-related ROI-deconvolution GLM

Figures 5 and 6 below show the results of deconvolution random effects GLM analysis, allowing the estimation of the unmixed response to each motion pattern. In general, response curves are very similar irrespective of the smoothing or not of the true trajectory of biological motion. Indeed, there are no significant differences between curves of response to the same motion type (global or local) in any ROI. We can observe that the curves in hMT+ seem to be anti-correlated in global and local conditions, and that the curves in pSTS seem biphasic, i.e., there seems to be an earlier and a later peak response.

**Figure 5:**
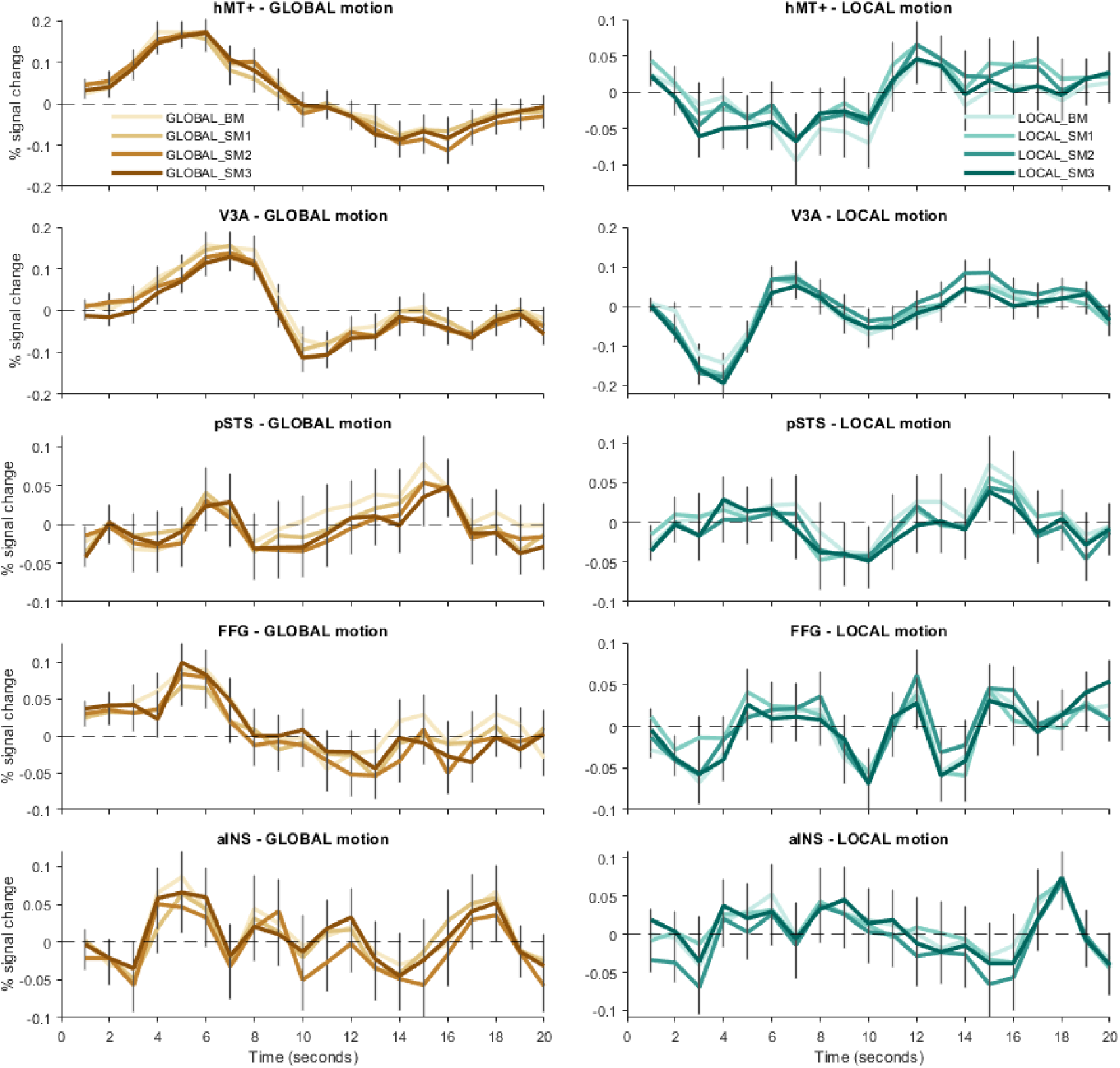
Deconvolution GLM plots of response to each type of motion event (global and local) in hMT+, V3A, posterior superior temporal sulcus (pSTS), fusiform gyrus (FFG) and anterior insula (aINS).

**Figure 6:**
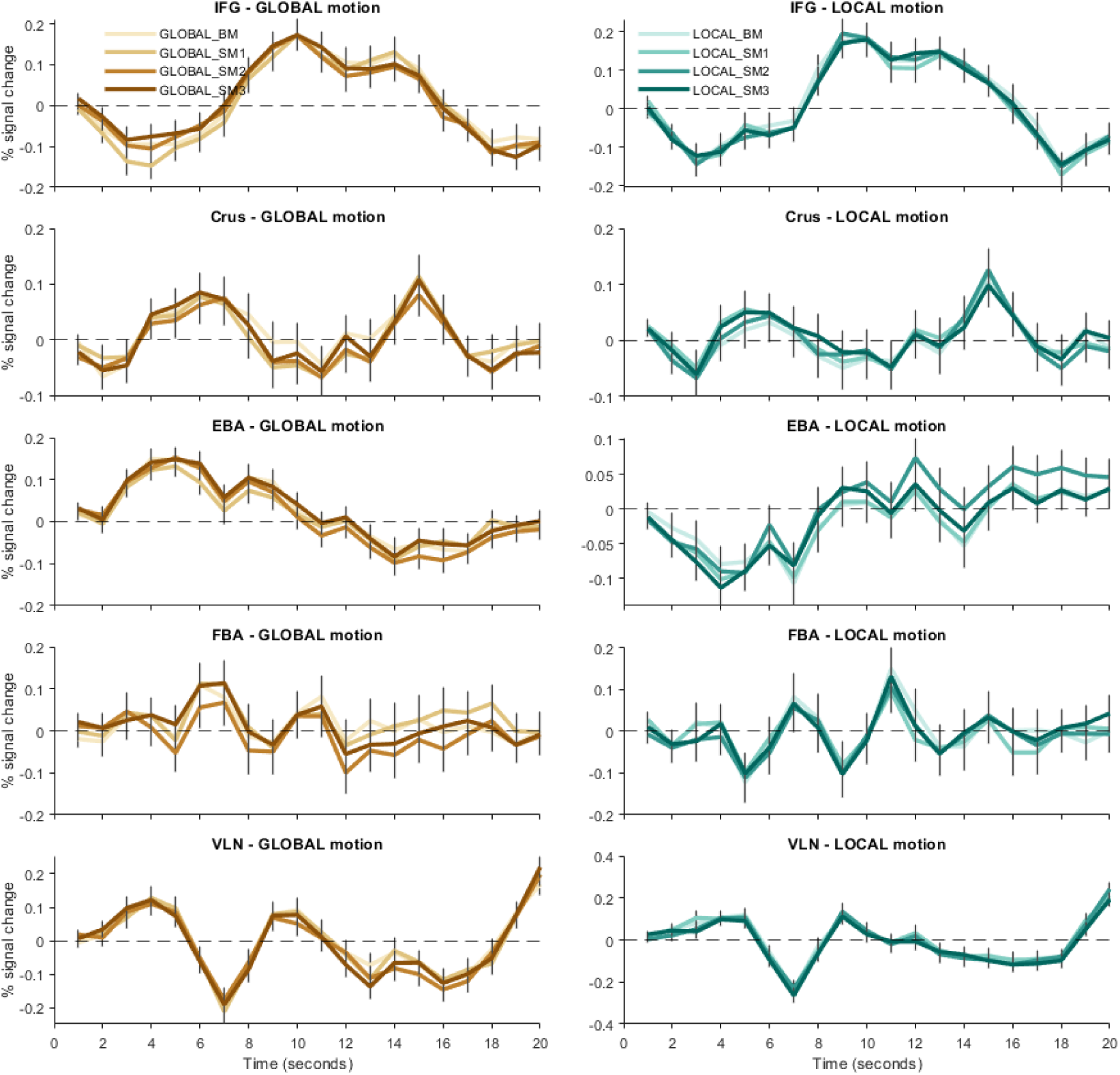
Deconvolution GLM plots of response to each type of motion event (global and local) in inferior frontal gyrus (IFG), lateral cerebellar lobule Crus I (Crus), extrastriate body area (EBA), fusiform body area (FBA) and ventral lateral nucleus (VLN) of the thalamus.

#### Representational Similarity Analysis (RSA)

### RSA of spatial activity patterns from univariate standard convoluted GLM

Figures 7 and 8 below show the RSA of spatial activity patterns of response to each type of motion, in each ROI. We can observe that spatial activity patterns of response in hMT+ and V3A, together with FFG, EBA and FBA, distinguish local from global motion type, as the four conditions of each type are well separated (clearly in 2D/3D MDS plots). Furthermore, these ROIs seem to also distinguish different motion profiles within motion type, to a certain degree. Although traditionally reported as the pivotal region for perception of biological motion, pSTS does not present this pattern of separated responses, which might represent a sort of invariance.

**Figure 7:**
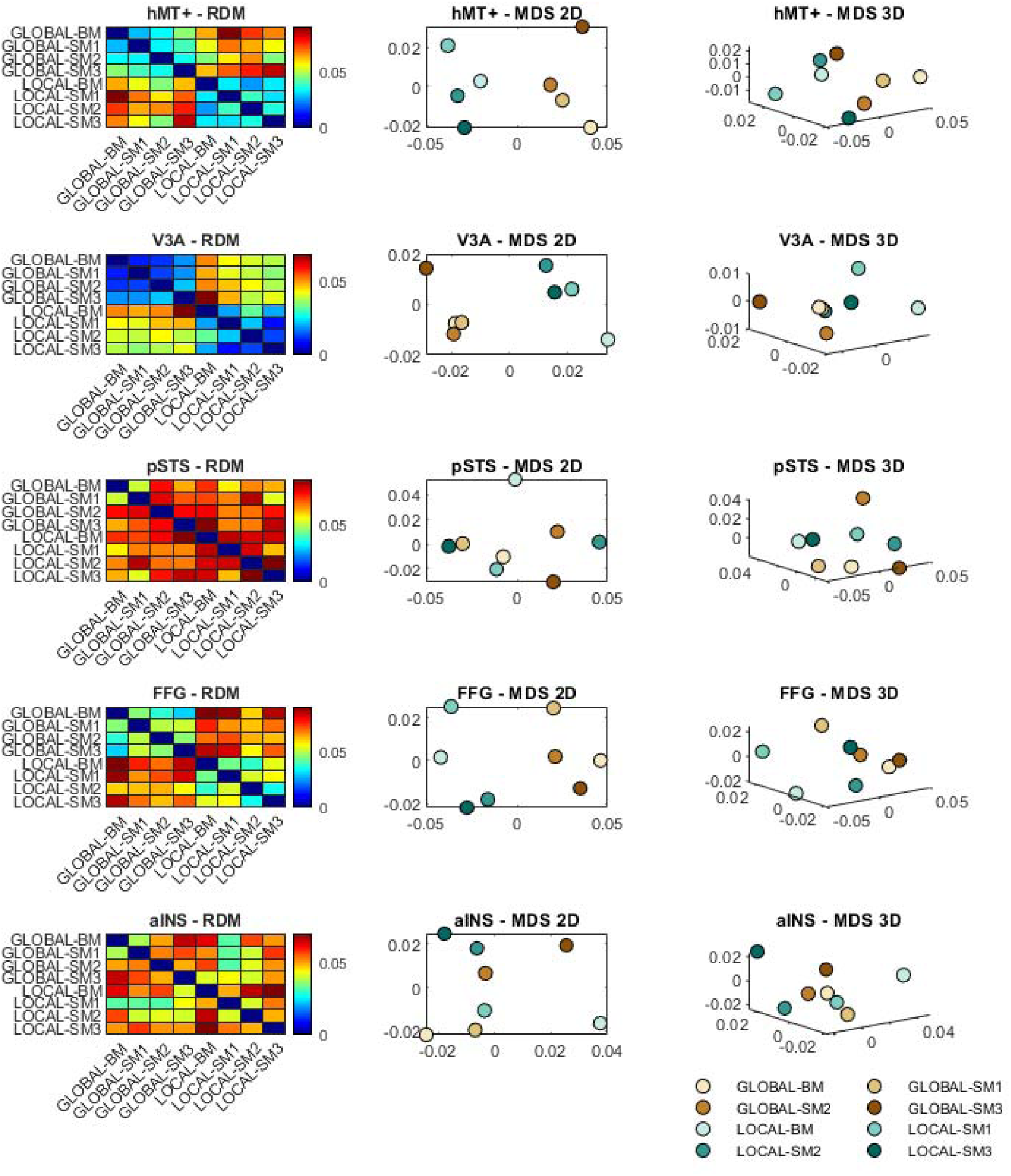
Representational similarity analysis of spatial patterns of response to each type of motion event (global and local) in hMT+, V3A, posterior superior temporal sulcus (pSTS), fusiform gyrus (FFG) and anterior insula (aINS). Each representational distance (or dissimilarity) matrix (RDM) is shown on the left, while 2D and 3D multi-dimensional scaling (MDS) plots are shown in the same row at the center and on the right, respectively. For each ROI the MDS plots show a point on the calculated 2D/3D coordinates as well as a label identifying the condition.

**Figure 8:**
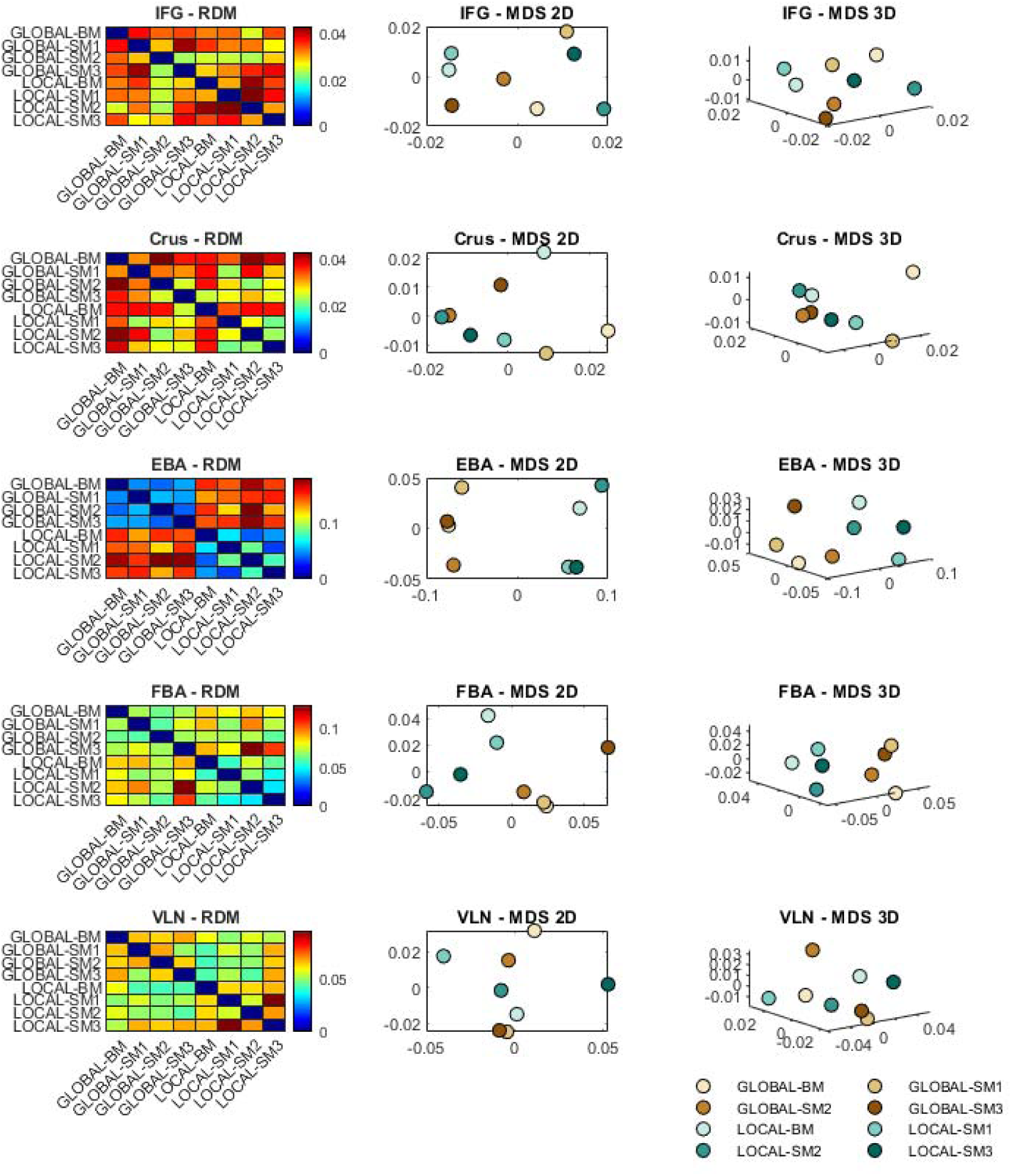
Representational similarity analysis of *spatial patterns of response* to each type of motion event (global and local) in in inferior frontal gyrus (IFG), lateral cerebellar lobule Crus I (Crus), extrastriate body area (EBA), fusiform body area (FBA) and ventral lateral nucleus (VLN) of the thalamus. Each representational distance (or dissimilarity) matrix (RDM) is shown on the left, while 2D and 3D multi-dimensional scaling (MDS) plots are shown in the same row at the center and on the right, respectively. For each ROI the MDS plots show a point on the calculated 2D/3D coordinates as well as a label identifying the condition.

### RSA of estimated HRF from deconvolution GLM

Figures 9 and 10 below show the RSA of temporal activity patterns of response to each type of motion, in each ROI. We can observe that temporal activity patterns of response distinguish local from global motion type, as the four conditions of each type are well separated, particularly in hMT+, V3A, FFG, EBA and FBA. We thus replicate the results of the spatial RSA from the same analysis perspective.

**Figure 9:**
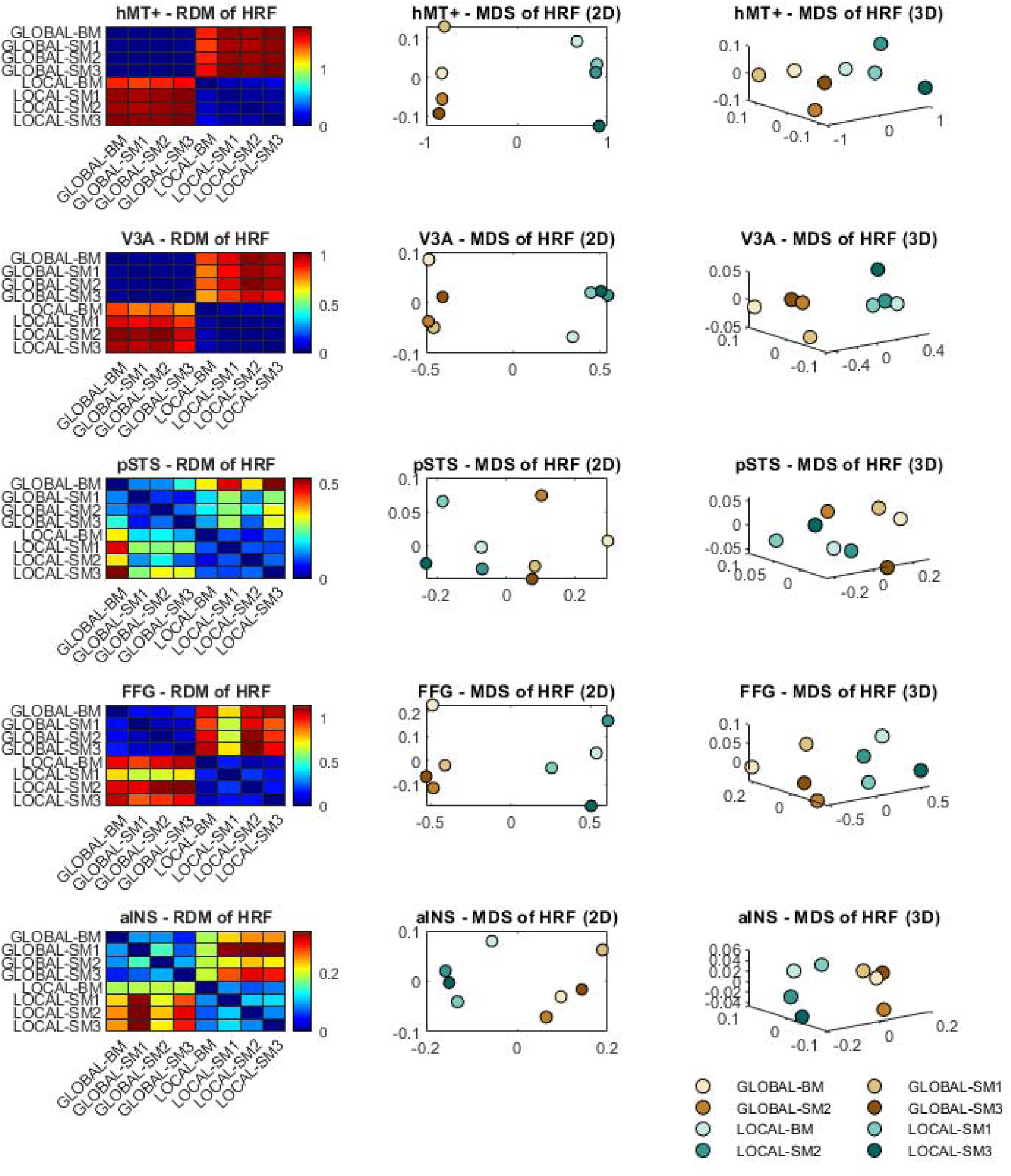
Representational similarity analysis of *temporal patterns of response* (the estimated HRF) to each type of motion event (global and local) in hMT+, V3A, posterior superior temporal sulcus (pSTS), fusiform gyrus (FFG) and anterior insula (aINS). Each representational distance (or dissimilarity) matrix (RDM) is shown on the left, while 2D and 3D multi-dimensional scaling (MDS) plots are shown in the same row at the center and on the right, respectively. For each ROI the MDS plots show a point on the calculated 2D/3D coordinates as well as a label identifying the condition.

**Figure 10:**
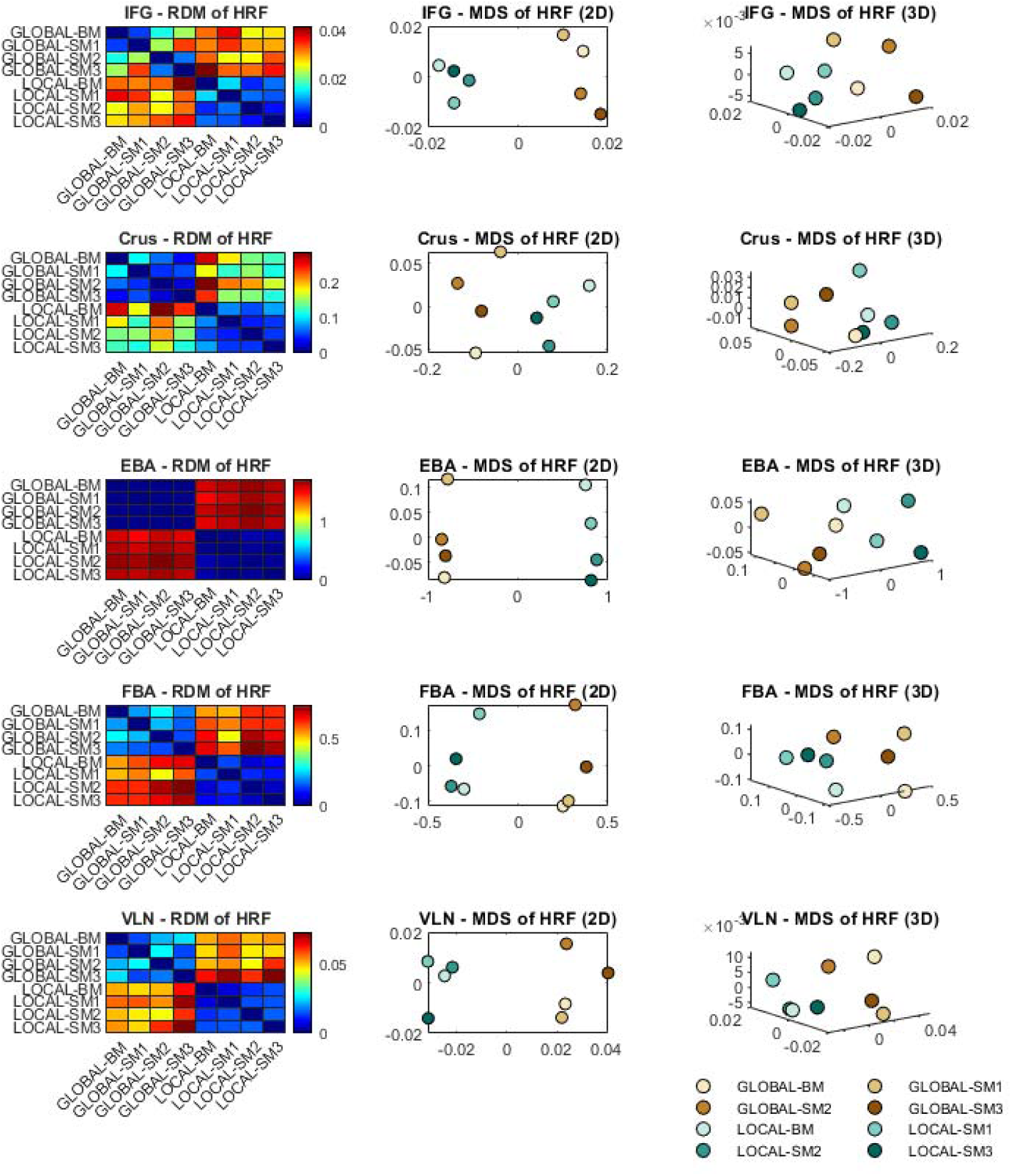
Representational similarity analysis of temporal patterns of response (the estimated HRF) to each type of motion event (global and local) in in inferior frontal gyrus (IFG), lateral cerebellar lobule Crus I (Crus), extrastriate body area (EBA), fusiform body area (FBA) and ventral lateral nucleus (VLN) of the thalamus. Each representational distance (or dissimilarity) matrix (RDM) is shown on the left, while 2D and 3D multi-dimensional scaling (MDS) plots are shown in the same row at the center and on the right, respectively. For each ROI the MDS plots show a point on the calculated 2D/3D coordinates as well as a label identifying the condition.

### Second-level RSA of activity patterns

Figure 11. below shows a second-level representational similarity analysis at the ROI level, i.e., the calculated similarity structure between conditions in each ROI is itself related to RDMs from every other ROI, producing the second-level RDM. This helps to understand the similarity of representational principles of brain regions. We can observe that hMT+ and V3A are closely together and clearly distant from the other regions, while EBA, FBA and FFG seem to be also separate from other higher-order regions in this representational space. On the other hand, the pSTS is closer to high-level regions and less well isolated from early-level regions.

## Discussion

In this study we used, for the first time, as to our knowledge, event-related deconvolution GLM analysis and representational similarity analysis (RSA) to study the neural correlates of the two-stage framework for neural processing of biological motion: life motion perception, using single dot stimuli that only contained local animacy properties, and global biological motion perception. We hypothesized that the first stage is involved in local and global motion, while the second stage is involved mainly in global motion processing. We found that the first stage relies on early visual areas hMT+ and V3A and higher-level areas FFG and EBA, which seem to discriminate local and global motion life motion. The second stage seems to rely on pSTS, which shows invariance to the subtype of biological motion.

**Figure 11:**
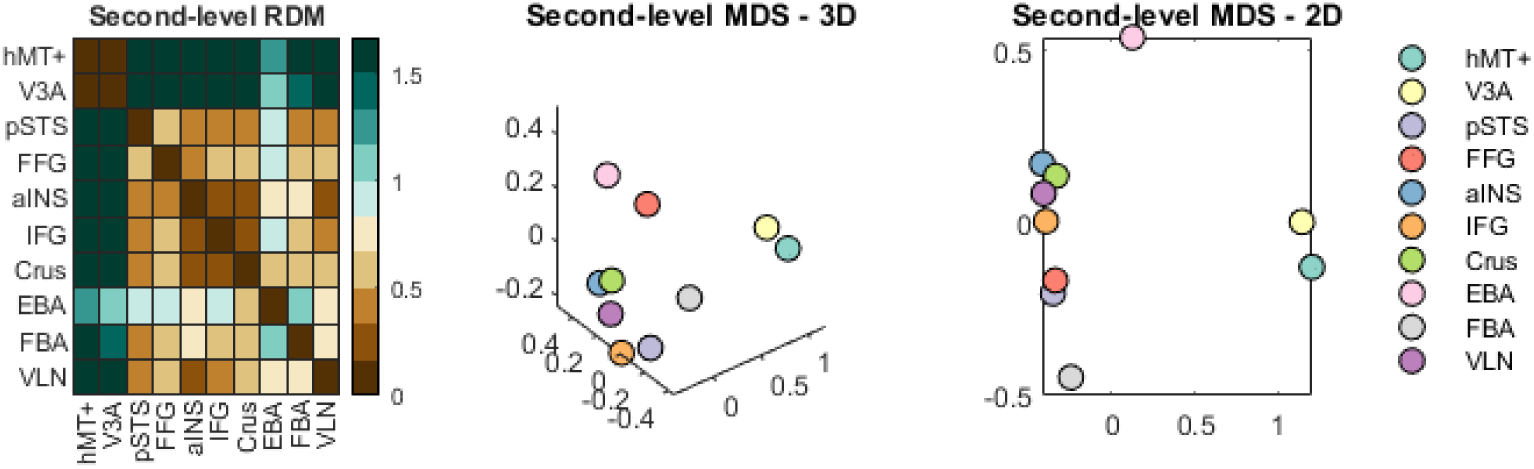
Second-level representational similarity analysis of spatial patterns of response to all types of motion (global and local) in each ROI. The representational distance (or dissimilarity) matrix (RDM) is shown on the left, while 2D and 3D multi-dimensional scaling (MDS) plots are shown at the center and on the right, respectively. The MDS plots show a point on the calculated 2D/3D coordinates as well as a label identifying the ROI.

Our observation that pSTS is involved in biological motion perception was expected, and adds to previous studies highlighting the pSTS as a key brain region for biological motion perception (Gilaie-Dotan et al., 2011), as well as gaze recognition, and animacy perception (Dasgupta et al., 2017; Hillebrandt et al., 2014). The observed invariance property is also consistent with this notion. Previous studies have addressed global forms of biological motion processing by using point-light displays (Johansson, 1973), which allow coherent articulated human motion to be integrated and perceived, despite the absence of form cues from the body (Troje, 2013). As such global patterns might also elicit high level cognitive processing, such as gender identification and even emotional processing, they have accordingly been used in studies of conditions characterized by social, behavioral, or emotional deficits (Yang et al., 2017). Several neuroimaging studies employing these kind of tasks and displays have investigated the neural processing of biological motion in in disorders of social cognition such as autism (Jack et al., 2017; Kaiser et al., 2010; Puglia and Morris, 2017). Both activation and connectivity patterns support the hypothesis that the pSTS serves as a hub of the social brain network (Alaerts et al., 2017; Dasgupta et al., 2017; Hillebrandt et al., 2014; Puglia and Morris, 2017).

However, it is known mainly from psychophysical data that humans (and also animals (Troje and Aust, 2013; Vallortigara and Regolin, 2006)) can readily judge direction of motion from inverted and scrambled point light displays if the dots associated with the local motion the feet remain intact (Troje, 2013, 2002; Troje and Westhoff, 2006). The authors interpret their findings as evidence for a visual filter that serves as a general detection system for articulated terrestrial animals: a local ‘life-detector’. However, the neural correlates of this life detector are still unknown. In this fMRI study we focused also on more basic aspects of biological motion processing, namely local biological motion in a perceptual decision task, which provides cues to discriminate animate from inanimate objects. Our stimulation paradigm using very short presentation of single dot stimuli free of structural cues elicited the recruitment of a neural system supporting the perceptual strength of the basic life motion detection (Hirai et al., 2011a).

Presenting briefly one dot representing a foot step or point light displays of a complete human body, while maintaining participants constantly engaged in perceiving biological motion, we expected to observe more localized activation in comparison to the network activated traditionally by global biological motion. Furthermore, we used deconvolution GLM to estimate responses to individual global and local motion patterns, and representational similarity analysis (RSA) to elucidate the similarity of brain regions regarding their co-activation patterns to each type of biological motion.

We observed a significant effect of ROI on percent BOLD signal change, with follow-up comparisons revealing that the response was overall higher in hMT+ than in all other ROIs. A previous meta-analysis (Grosbras et al., 2012) had already suggested that hMT+ is activated by biological motion but stated that previous studies were not able to identify whether such responses can be specifically linked to human movement perception, given the nature of used contrasts. Our result highlights the role of this region in early level perceptual decision involving biological motion, which is compatible with predictive coding accounts of visual motion prediction (Vetter et al., 2015), and do indeed suggest that hMT+ indeed plays a role in tasks requiring discrimination of true life motion. This further corroborates previous evidence that hMT+ activity reflects the complex motion pattern present in biological motion stimuli, while that of STS reflects the action portrayed in the stimuli (Peuskens et al., 2005). In fact, hMT+ has been implicated in altered activity and connectivity patterns during perception of biological motion both in health (Hillebrandt et al., 2014) and disease (Herrington et al., 2007). The signal in pSTS, commonly associated with biological context processing, was significantly lower also than in FFG and EBA, the former showing a response significantly higher than in all other ROIs except hMT+, FFG and V3A. Notably, we also found an interaction between ROI and type of motion (global or local), indicating that the differences in brain responses to different types of motion are dependent on the ROI considered. We highlight that although the amplitude of responses is higher for global motion relative to local motion, the sub-network of preferentially activated regions is the same in both situations, including hMT+, V3A, EBA and FFG. A recent study with a single dot evoking controlled degrees of perceived animacy found BOLD signals reflecting perceived animacy in one intraparietal region (IPS) (Schultz and Bülthoff, 2019). However, their stimuli did not involve perception of human action and thus may not engage ventral or lateral temporal regions such as we found. Chang and colleagues have also demonstrated the involvement of the hMT+, V3A and EBA in perception of global and distributed local biological motion cues (Chang et al., 2018). Our study further highlights the involvement of FFG as part of the first stage in biological motion processing. While Sokolov and colleagues have shown that FFG is also connected to higher-order regions, indicating a parallel pathway for biological motion perception (Sokolov et al., 2018), they have used only stimuli with global biological motion, suggesting the role of the FFG in high-level socio-cognitive processing and social information integration, which have been observed to be altered in autism spectrum disorder (Yang et al., 2017). Our results suggest that these regions are the core supporting the first stage of global and also local biological motion perception. This is of obvious evolutionary value, as it shows that this core contains information on the generalizing invariant encoding the presence of a living being, irrespective of its shape. Such sensitivity to life motion biomechanical details places these regions in the initial - first stage -biological motion processing network.

On the other hand, the response profile in pSTS suggests that this region supports a second stage in biological motion processing, irrespective of the context containing local or also global information, which defines invariance. This is in accordance with previous studies revealing significant connectivity patterns of lower-level regions with pSTS, as well as the cerebellum (Jack et al., 2017; Sokolov et al., 2018). By isolating the intrinsic motion component from other high-level sources of information about actor and action we were able to portray an early local computation (first stage) and a second stage of biological motion perception and to identify its neural correlates. There is previous evidence that action perception involves at least two parallel pathways, relying on complementing but dissociable functions of EBA and pSTS, which carry information about body for and motion, respectively (Vangeneugden et al., 2014). However, local biological motion was not addressed in this study.

Our results are in accordance with recent evidence of the existence of an hierarchical third visual pathway on the lateral brain surface, that is anatomically segregated from the dorsal and ventral pathways, supporting biological motion (Pitcher and Ungerleider, 2020). This pathway begins in primary visual cortex and projects into pSTS via hMT+ (Gschwind et al., 2012; Pitcher et al., 2020). Furthermore it includes EBA, which can overlap with hMT+ (Downing et al., 2007), and responds more to moving than static bodies (Pitcher et al., 2019). In addition, the temporoparietal junction (TPJ) (an adjacent brain area anterior and superior to the STS) responds to theory of mind tasks, in which participants are required to interpret the actions of characters from motion patterns, and its activity has been shown to be altered also in contexts of social deficits (Madeira et al., 2021). The findings of our study suggest the differentiation of this third visual pathway in two processing stages, culminating in pSTS, and the addition of the FFG to the first stage together with hMT+, V3A and EBA. However, the different representational activation patterns in these regions in response to local vs. global motion, clearly revealed by RSA (see MDS plots), indicate that there is a dichotomic contribution of the traditional dorsal stream (hMT+ and V3) and ventral stream (EBA and FFG) regions in the early processing of biological motion. The core supporting the first-stage clearly separates local from biological motion, while pSTS does not. This is evident in the second-level RSA, in which we can observe that these four regions are separated from the remaining in the 2D and 3D MDS plots, indicating that their activation patterns are more similar between them than the other regions, while pSTS is somehow less distinguishable from other higher-level regions.

We cannot exclude that life motion perception, like in other perceptual decision studies, might be modulated by top-down mechanisms originated in higher level-regions such as the insula (Duarte et al., 2017; Rebola et al., 2012; Stottinger et al., 2015). The suggestive involvement of the cerebellum, observed in the response profiles, is not unexpected considering it outputs to premotor cortex or supplementary motor area and is involved in sensory guidance and planning of movements based on external cues. Similarly, the ventral lateral nucleus (VLN) holds extensive projections to the motor and premotor cortex (Kievit and Kuypers, 1975; Shinoda et al., 1993) and receives considerable afferents from the cerebellum (dentate nucleus) and the globus pallidus (Carpenter and Strominger, 1967). As such, the VLN is classically thought to act as a major relay for motor planning and coordination (e.g., (Daskalakis et al., 2004)) and has been shown to be sensitive to both form and kinematic information in biological motion (Chang et al., 2018). Although our results are not conclusive of the involvement of this (primarily motoric) thalamic structure for biological motion perception, the RSA of temporal response patterns reveals the need to consider the possibility of a much earlier (pre-cortical) encoding both local and global biological motion. This is in accordance with the implication of the cerebellum in the processing of biological motion (Sokolov et al., 2012), yet requires further validation with more detailed investigation.

In conclusion, by studying local life motion perception, devoid of global shape, coherent articulated motion, social cues and other sources of information, we were able to segregate two processing stages of biological motion perception. Given that human adults have very considerable experience of the visual world it is not surprising if they have acquired the perceptual skill necessary for extracting the most informative features of biological motion from a single dot representing a real step (joint) trajectory (Peelen et al., 2006). The two-stage framework for neural processing of biological motion supports the idea of an evolutionarily ancient neural mechanism that is activated by the perception of local and global biological motion in humans.

## Funding

This work was supported by the Portuguese Foundation for Science and Technology (grant numbers UID/04950B/2020, UID/04950P/2020, DSAIPA/DS/0041/2020, PTDC/PSI-GER/30852/2017, PTDC/PSI-GER/1326/2020 and PTDC/MEC-NEU/31973/2017). FCT also funded an individual grant to JVD (Individual Scientific Employment Stimulus 2017 - CEECIND/00581/2017).

## Acknowledgments

We would like to thank the participants for their involvement in this study. We are also very grateful to Sónia Afonso and Tânia Lopes for the help with MRI setup and scanning.

## Competing interests

The authors declare that no competing interests exist.

## Declarations

The data and the code used in the study are available from the corresponding author upon reasonable request. Data and code sharing must comply with the requirements of the University of Coimbra and with approval by the ethics board.

## Supplementary material

### Functional localization of hMT+ and V3A

The average number of voxels and MNI coordinates of functionally localized left and right hMT+ and V3A are shown in Table S1.

**Table S1.**
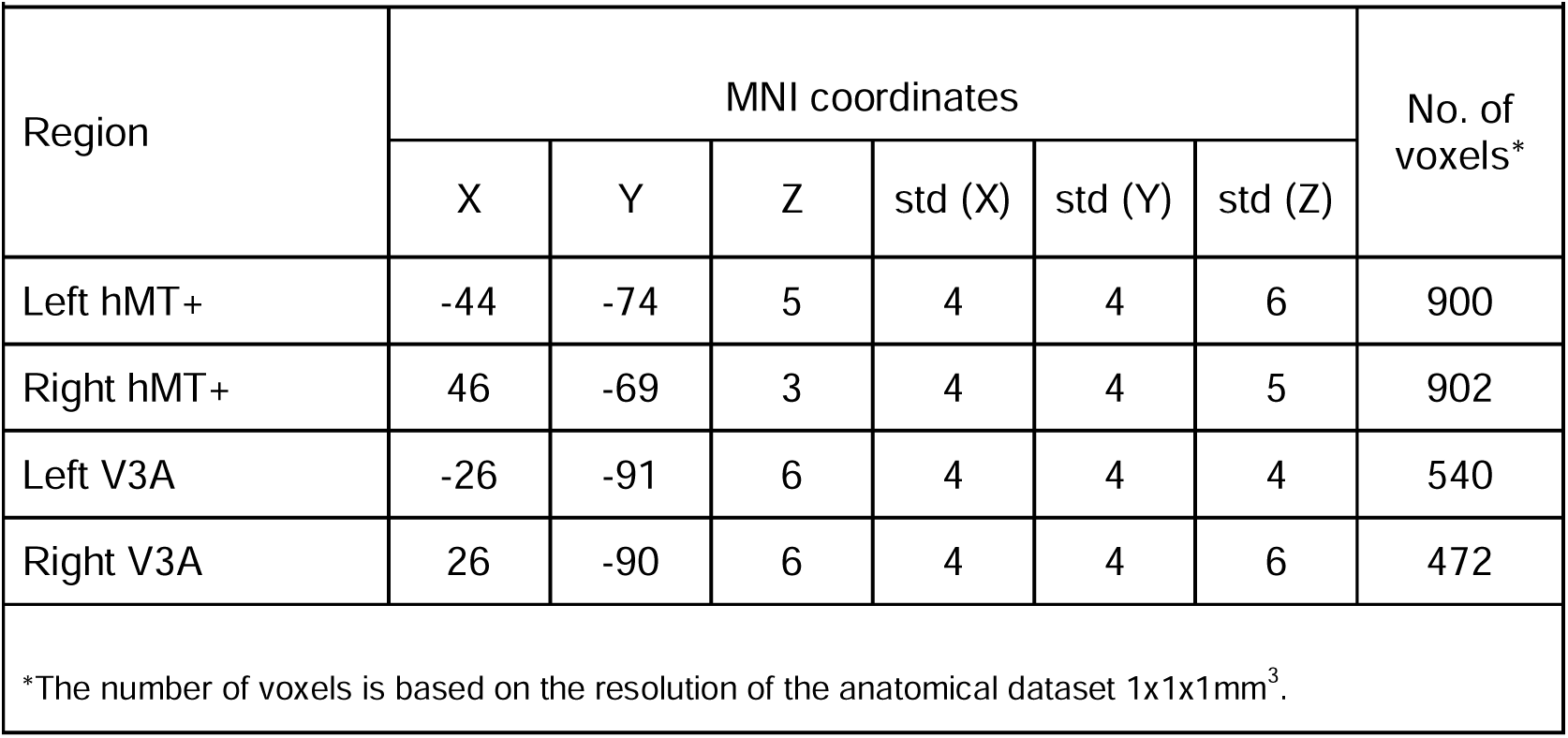
Summary of the individual localization of hMT+ and V3A regions-of-interest [p-value(Bonf)<0.05].

## Notes

### Competing Interest Statement

The authors have declared no competing interest.

