## Supplementary Table S1 for "A two-stage framework for neural processing of biological motion"

### Functional localization of hMT+ and V3A

The average number of voxels and MNI coordinates of functionally localized left and right hMT+ and V3A are shown in Table S1.

| **Table S1. Summary of the individual localization of hMT+ and V3A regions-of-interest [p-value(Bonf)<0.05]** | | | | | | | |
| --- | --- | --- | --- | --- | --- | --- | --- |
| Region | MNI coordinates | | | | | | No. of voxels* |
|  | X | Y | Z | std (X) | std (Y) | std (Z) |  |
| Left hMT+ | -44 | -74 | 5 | 4 | 4 | 6 | 900 |
| Right hMT+ | 46 | -69 | 3 | 4 | 4 | 5 | 902 |
| Left V3A | -26 | -91 | 6 | 4 | 4 | 4 | 540 |
| Right V3A | 26 | -90 | 6 | 4 | 4 | 6 | 472 |
| *The number of voxels is based on the resolution of the anatomical dataset 1x1x1mm^3^. | | | | | | | |
